# CX_3_CR1 Fate-Mapping In Vivo Distinguishes Cochlear Resident and Recruited Macrophages After Acoustic Trauma

**DOI:** 10.1101/2025.08.04.668473

**Authors:** Sree Varshini Murali, Andrew R Stothert, Elyssa Pereyra, Lyudmila V Batalkina, Tejbeer Kaur

## Abstract

Cochlear injury activates the resident macrophages (RM) and recruits the blood-circulating monocytes and monocyte-derived macrophages (Mo/Mo-M), but their specific functions in the injured cochlea are unknown. It is well established that the chemokine fractalkine receptor (CX_3_CR1), expressed by cochlear macrophages, influences the density of those macrophages and promotes synaptic repair and spiral ganglion neuron survival in the injured cochlea. As CX_3_CR1 is expressed on both RM and Mo/Mo-M, it remains unclear if CX_3_CR1-expressing RM and Mo/Mo-M are distinct and differentially promote SGN survival after cochlear injury. Here, we demonstrate the use of fate mapping via a tamoxifen-inducible CX_3_CR1 mouse model (CX_3_CR1^YFP−CreERT2/wildtype^:R26RFP) wherein CX_3_CR1-expressing RM and Mo/Mo-M are endogenously labeled with different fluorescent reporters to define the heterogeneity in cochlear macrophages regarding their origin, turnover, spatiotemporal distribution, morphology, and fate following a loud acoustic trauma. After 60 days of tamoxifen injections at 4 weeks of age, long-lived cochlear RM were YFP+ RFP+ with 98.0 ± 1.7% recombinant efficiency, and short-lived blood-circulating CX_3_CR1 lineage (Mo/Mo-M) were YFP+ RFP-with 2.5 ± 1.1% recombinant efficiency. Following an acoustic trauma of 112 dB SPL at 8-16 kHz octave band for 2 hours, morphologically similar RM and Mo/Mo-M were observed in the spiral ganglion, lamina, ligament, and around the sensory epithelium. Quantification of RM and Mo/Mo-M in the spiral lamina and ganglion revealed distinct spatial and temporal distribution patterns. Furthermore, recruited Mo/Mo-M expressed classical monocyte markers such as Ly6C and CCR2. Both RM and Mo/Mo-M were positive for proliferation marker, Ki67, and negative for apoptotic marker, cleaved caspase-3, suggesting that the overall increase in macrophage numbers in the noise-injured cochlea is a contribution of both the proliferation of RM and recruitment of Mo/Mo-M. Probing for blood-clotting protein, fibrinogen, showed its presence in the cochlea after acoustic trauma, suggesting vascular damage that positively and strongly correlated with the time course of recruitment of blood-circulating Mo/Mo-M in the noise-injured cochlea. These data imply that macrophages in the noise-injured cochlea are heterogeneous regarding their ontogeny, distribution, and fate. They offer a robust tool to study the precise roles of resident and recruited macrophages in healthy and pathological ears.

**Summary:** Using the novel CX_3_CR1 fate-mapping model, our data uncover diversity of macrophages with respect to their ontogeny, turnover, spatiotemporal distribution, and fate in the normal and noise-injured cochlea.

## BACKGROUND

The mononuclear phagocyte system represents a subgroup of leukocytes or white blood cells originally described as a population of bone marrow-derived myeloid cells that circulate in the blood and spleen as monocytes and populate tissues as macrophages in the steady state and during inflammation **(Geissmann et al., 2010, van Furth and Cohn, 1968).** Monocytes are innate-immune effector cells that migrate from blood to tissues during injury or infection and can also differentiate into macrophages during inflammation **(Ginhoux and Jung, 2014, Jung, 2018).** Macrophages are tissue-resident phagocytic cells that play a central role in both steady-state tissue homeostasis, repair, and injury. Macrophages are involved in phagocytosis of apoptotic cells, production of growth factors, pro-inflammatory and anti-inflammatory cytokines to promote wound repair during injury, and processing and presenting antigens to T-lymphoid cells **(Ginhoux and Jung, 2014)**. The resident macrophages in adult tissue in the steady state might arise only from yolk sac-embryonic macrophages (brain microglia), from both yolk sac-embryonic macrophages and fetal liver monocytes (Langerhans cells in skin, alveolar macrophages in lung and Kupffer cells in liver) and can also be replenished by adult bone marrow-derived monocytes (heart and gut macrophages) **(Ginhoux and Guilliams, 2016)**. Hence, macrophages are a heterogeneous cell type that respond differently based on the environmental niche, anatomical location, and distinctive origin.

It is well-established that the developing and adult mammalian cochlea contains a population of local or resident macrophages that are distributed in the spiral ganglion, osseous spiral lamina, spiral limbus, spiral ligament, and stria vascularis of the lateral wall **(Dong et al., 2018, Hirose et al., 2005, Hosoya et al., 2023, Liu et al., 2018, O’MalleyNadol and McKenna, 2016, Okano et al., 2008, Shi, 2010)**. Mild to loud noise trauma, aminoglycoside- or cisplatin-induced ototoxicity, infection, or normal aging of the cochlea is associated with robust activation of resident macrophages and an overall increase in the number of macrophages **(Frye et al., 2017, Hirose et al., 2005, Kaur et al., 2015, Sung et al., 2024, Yang et al., 2015)** Whether this increase in macrophage numbers in the damaged cochlea is due to local proliferation of resident macrophages (RM) or recruitment of monocytes (Mo) from blood circulation and their differentiation into macrophages (Mo-M) or both, remains ambiguous. We have recently demonstrated that macrophages play a vital protective role in the damaged cochlea by promoting the long-term survival of spiral ganglion neurons (SGN) and repair of damaged ribbon synapses via neuron-immune fractalkine (CX_3_CL1-CX_3_CR1) signaling between SGNs (which express CX_3_CL1 ligand) and macrophages (which express CX_3_CR1 receptor) **(Kaur et al., 2019, KaurOhlemiller and Warchol, 2018, Kaur et al., 2015, Manickam et al., 2023, Manickam et al., 2024)**. Notably, CX_3_CR1 is present on both RM and Mo/Mo-M and is morphologically similar **(Jung et al., 2000)**, making these two macrophage populations indistinguishable by standard immunohistochemical techniques. Thus, it remains unclear whether CX_3_CR1-expressing RM or infiltrated Mo/Mo-M are functionally distinct and differentially promote SGN survival in the injured cochlea. To define the mediators of macrophage-induced neuroprotection and to harness the protective capacity of these cells clinically, it is necessary to delineate the specific cell type (i.e., CX_3_CR1-expressing RM or Mo/Mo-M) that promotes synaptic repair and SGN survival after cochlear injury.

In the current study, we utilized a fate mapping technique by crossing a tamoxifen-inducible Cre mouse line (CX_3_CR1YFP−CreER/YFP−CreER) to a red fluorescence protein (RFP) Cre reporter mouse line (R26RFP) **(Goldmann et al., 2013, O’KorenMathew and Saban, 2016b, Parkhurst et al., 2013, Plemel et al., 2020a)** wherein CX_3_CR1-expressing RM and Mo/Mo-M are endogenously labeled with different fluorescent reporters to define the heterogeneity in cochlear macrophages regarding their origin, spatiotemporal distribution, morphology, and fate following an acoustic trauma that causes permanent hearing loss. We show here that the progeny of this crossing, CX_3_CR1YFP−CreER/wt:R26RFP, can be used to differentially label resident macrophages (RM) versus recruited Mo-derived cells (Mo/Mo-M) in the cochlea. This system works by tamoxifen pulsing mice to temporarily activate Cre in constitutively CX_3_CR1-positive (YFP+) cells, i.e., Mo and cochlear resident macrophages. The activated Cre removes a stop codon controlling RFP expression such that YFP+ cells become RFP+. Once the tamoxifen is no longer bioavailable, Cre is inactive, and no new YFP+ cells can express RFP. This step is followed by a “wash out” period of several weeks wherein RFP expression is lost in Mo (as they turnover in circulation because of ongoing, lifelong hematopoiesis), whereas expression is naturally and indefinitely retained by cochlear RM (because they are long-lived).

Using this fate mapping system, in conjunction with flow cytometry and histology, we demonstrate that macrophages in the noise-injured cochlea are heterogeneous regarding their ontogeny, spatial and temporal distribution, and fate. Furthermore, the overall increase in macrophages in the noise-injured cochlea is due to the contribution of both the proliferation of RM and the recruitment of Mo/Mo-M, the latter is associated with a leaky vasculature. This study offers a robust tool to study the isolated roles of resident and recruited macrophages in healthy and pathological ears and to define their neuroprotective functions in the noise-injured cochlea. These findings are likely relevant in other cochlear disease models with myeloid cell involvement, such as hearing loss due to ototoxic drugs, bacterial and viral infections, synaptopathy, biological aging, genetics, and sudden sensorineural hearing loss, as well as associated comorbidities including diabetes, cardiovascular diseases, depression, dementia, and cognitive decline.

## MATERIALS AND METHODS

### Mice

CX_3_CR1YFP−CreER/YFP−CreER [stock No. 021160] and the Cre reporter mouse line, Rosa-lsl-tdTomato (R26RFP) [stock No. 007914], were purchased from Jackson Laboratories (Bar Harbor, Maine). Efforts were made to minimize animal suffering and reduce the number of animals used for experiments. The animals were housed in a temperature- and humidity-controlled environment in autoclaved cages in groups of 5 under a 12 h light/12 h dark cycle and fed *ad libitum*. To confirm the genotype of the mice, extracted genomic DNA was subjected to polymerase chain reaction (PCR) using forward and reverse primers described in **Table 1**. All aspects of animal care, procedures, and treatment were conducted according to the National Institute of Health guidelines and approved by the Animal Care and Use Committee of Creighton University and Rutgers University.

**Table 1.**
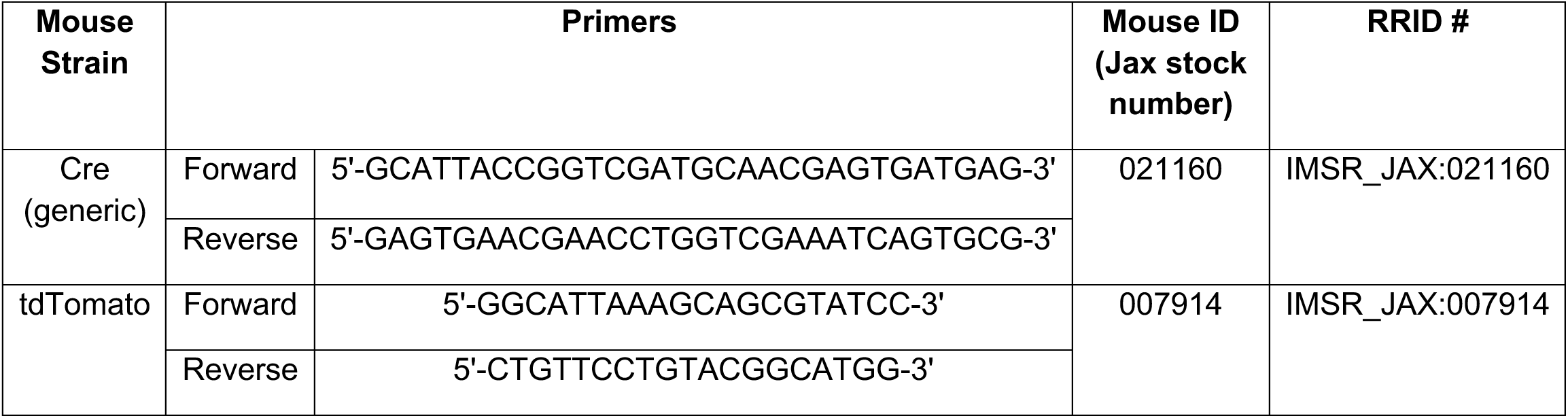
List of primers to genotype mice by PCR.

### Tamoxifen or corn oil (vehicle) injection

Tamoxifen (Sigma-Aldrich) was dissolved in corn oil (Sigma-Aldrich) to a stock concentration of 10 mg/ml. 75 mg/kg of tamoxifen was injected intraperitoneally (i.p.) twice, with one day in between injections. Mice were approximately 4 weeks of age when given tamoxifen.

### Noise exposure

For noise exposure, we implemented the methodology described in our previous work **(Manickam et al., 2023)**. Briefly, conscious and freely moving mice were subjected to a 2-hour exposure to octave band noise (8–16 kHz) at a consistent intensity of 112 dB SPL, which imparts permanent hearing loss **(Sautter et al., 2006)**. The exposure was performed within a sound-attenuating chamber (WhisperRoom) lined with acoustic foam. The mice were housed either singly or in pairs within modified compartmentalized cages from which food, water, and bedding materials had been removed. These cages were arranged with a maximum of two units positioned directly beneath an exponential horn attached to a speaker (JBL). The acoustic stimulus was computer-generated via customized LabVIEW software interfaced with an audio card (Lynx E22), producing a filtered pure tone (8-16 kHz) that was subsequently amplified via a power amplifier (Crown Audio) connected to the horn speaker. Each exposure session was preceded by calibration via a quarter-inch condenser microphone (PCB) to verify the target sound pressure level (SPL), which varied by ±1 dB across the compartments in the cage. The control groups consisted of unexposed age-matched mice.

### Cochlea and Blood harvest

Mice were deeply anesthetized with lethal doses (250 mg/kg) of pentobarbital sodium (Trade Name-Fatal Plus) or pentobarbital sodium and phenytoin sodium (trade name-Euthasol). Before respiratory arrest, mice were perfused by transcardiac route with phosphate-buffered saline (PBS) (Fisher Scientific, catalog #BP661-10) or 4% paraformaldehyde (PFA) (Fisher Scientific, catalog #50980495) in 0.1 M phosphate-buffered (PB) solution. Temporal bones and blood were harvested. Blood was collected via terminal cardiac puncture before perfusion. Withdrawn blood was immediately incubated for 15 min at room temperature in a 0.2% Heparin solution (ScienCell, catalog #0863) to prevent coagulation. Heparinized blood (1 ml) was treated with 10 ml of 1× Red Blood Cell Lysis Buffer (Invitrogen, catalog #00-4333-57) for 5 min at room temperature with gentle agitation and subsequently thoroughly washed with PBS, pelleted at 300 × *g* for 5 min, and then resuspended in 1 ml of cell staining buffer (BioLegend, catalog #420201) before cell counting, immunolabeling, and flow cytometry.

### Flow cytometry

Single-cell suspensions of lysed blood were incubated for 10 min on ice in a blocking solution containing 2% Fc Block and subsequently stained with a combination of fluorophore-conjugated primary antibodies against CD45, CD11b, Ly6G, and Live/Dead Fixable Violet Dead Cell Stain. Samples were incubated in the dark and on ice for 30 min. After completion of staining, cells were washed with 1 ml of cell staining buffer (BioLegend, catalog #420201), pelleted at 300 × *g* for 5 min, resuspended in 200 μl of cell staining buffer, and fixed with 4% PFA in PBS. Data were acquired with the YETI Cell Analyzer (Propel Labs) using Everest Software (Bio-Rad Laboratories). Gating strategy and raw flow cytometry data were analyzed using FlowJo software (TreeStar). Cell gating was initially set using forward-scatter versus side-scatter plots of all events that corresponded to the size of cells. These cells were gated and labeled “blood cells.” These cells were then gated using the Live/Dead stain, gating for live cells only. Live cells were then gated for singlets (single-cell events) by comparing forward-scatter width and side-scatter area. Singlets were then analyzed for the presence of leukocytes using CD45 expression. CD45+ cells were then further analyzed for myeloid lineage using CD11b. Granulocytes were excluded by gating for CD45+ CD11b+ Ly6G+ cells. Finally, CD45+CD11b+Ly6G-myeloid cells in blood were analyzed for the percentage of RFP+ and RFP-expression at various days after tamoxifen injections. **Table 2** describes all antibodies and probes used in the study.

**Table 2.** List of antibodies and probes.

| Probes and antibodies | Host species | Isotype | Clone | Dilution/ volume | Vendor | Catalog # | RRID # |
| --- | --- | --- | --- | --- | --- | --- | --- |
| Anti-Mouse CD16/CD32 (Fc block) | Rat | IgG2bk | 2.4G2 | 2 µL/ 100 µL of cell suspension | BD Biosciences | 553142 | AB_394657 |
| Live/Dead Stain | NA | NA | NA | 1µL of 1:100 dilution | Invitrogen | L34967 | NA |
| APC-CD45 | Rat | IgG2bk | 30-F11 | 1:5 | BD Biosciences | 559864 | AB_398672 |
| BV711-CD11b | Rat | IgG2b | M1/70 | 1:5 | BioLegend | 101241 | AB_11218791 |
| PECy7-Ly6G | Rat | IgG2b | RB6-8C5 | 1:2 | Thermo Fisher Scientific | 25-5931-81 | AB_469662 |
| APC-CCR2 | Rat | IgG2b | 475301 | 1:100 | R&D Systems | FAB5538A | AB_10645617 |
| anti-CD45 | Goat | IgG | NA | 1:50 | R&D Systems | AF114 | AB_442146 |
| anti-GFP/YFP | Rabbit | IgG | NA | 1:500 | Thermo Fisher Scientific | A-11122 | AB_221569 |
| anti-GFP/YFP | Chicken |  | NA | 1:500 | Aves lab | GFP-1010 | AB_2307313 |
| anti-Cleaved Caspase 3 | Rabbit | IgG | NA | 1:400 | Cell Signaling | 9661S | AB_2341188 |
| anti-Ki67 | Rabbit | IgG | NA | 1:200 | Abcam | ab15580 | AB_443209 |
| anti-Fibrinogen | Rabbit | IgG | NA | 1:500 | Dako, Agilent | A0080 | AB_2894406 |
| anti-Ly6C | Mouse | IgG1 | NA | 1:100 | Santa Cruz Biotechnology | sc-271811 | AB_10707825 |
| anti-CCR2 | Rabbit | IgG | NA | 1:100 | Abcam | ab216863 | AB_2832204 |

954 (NA – not applicable)

### Immunohistochemistry

Cochlear microdissected whole mounts or frozen mid-modiolar cross sections (20-25 µm) were rinsed with PBS (Fisher Scientific, catalog #BP661-10) at least 3 times for 15 minutes each time and incubated at room temperature for 2 h in blocking solution containing 5% normal horse serum (Sigma-Aldrich, catalog #H0146) in 0.2% Triton X-100 (Sigma-Aldrich, catalog #A16046AP) in PBS. Tissue was incubated overnight at room temperature with the following primary antibodies: anti-CD45, anti-GFP/YFP (to enhance visualization of CX_3_CR1YFP-expressing macrophages), anti-cleaved caspase-3, anti-Ki67, anti-fibrinogen, anti-Ly6C, and anti-CCR2. Following incubation in primary antibodies, specimens were rinsed for 15 minutes in PBS, repeated 3 times, and treated for 2 h at room temperature in species-specific secondary antibodies conjugated to either DyLight-405 or -647 (1:500, Jackson ImmunoResearch Laboratories) or AlexaFluor-488, -546, -555, or -647 (1:500; Invitrogen). Tissue was rinsed for 15 minutes in PBS, repeated 3 times, and mounted in glycerol: PBS (9:1) and cover-slipped before confocal imaging. **Table 2** describes all antibodies and probes used in the study.

### Confocal imaging

Three- or four-color fluorescence imaging was performed using a Zeiss LSM 700 or 900 laser scanning confocal microscope (Carl Zeiss Microscopy). *Z*-series images using 5x, 10×, 20×, 40×, or 63× objectives were obtained. Image processing and quantitative analyses were performed using IMARIS (Oxford Instruments), Volocity 3D image (Quorum Technologies Inc.), and Image J (NIH).

### Resident and recruited macrophage counts

Resident macrophages (YFP+RFP+) and recruited monocytes and monocyte-derived macrophages (YFP+RFP-) were quantified in the spiral ganglion and the osseous spiral lamina (OSL) per cochlear section. Five-six mid-modiolar cochlear frozen sections, each 20 µm thick, were collected from sham-exposed and noise-exposed mice across eight time points: 0 hour (immediately), and 1, 3, 5, 7, 10, 15, and 30 days after noise exposure. Sections were immunolabeled for anti-GFP and anti-CD45 antibodies. Cells expressing YFP/GFP, RFP/tdTomato, and CD45 were quantified using IMARIS software and confirmed manually from 20x confocal images. Raw macrophage counts were reported for the OSL, while counts within spiral ganglion were normalized to the area of Rosenthal’s canal for apex, middle, and base cochlear region and expressed as macrophage density per 1000 µm².

### Fibrinogen Intensity measurement

Cochlear cryosections were immunolabeled for fibrinogen and imaged using an LSM 900 confocal microscope. The mean fluorescent intensity (MFI) was measured using the Image J software (NIH). A region of interest (ROI) was drawn around the site of fibrinogen signal, within Rosenthal’s canal, lower spiral ligament, and spiral limbus. The area and mean fluorescence of the ROI were measured. Similarly, ROI was drawn over the region with no fibrinogen signal within the Rosenthal canal, lower spiral ligament, and spiral limbus to measure the background. The final fibrinogen fluorescent intensity is reported as the average MFI of the ROI subtracted from the average MFI of the background, keeping the area constant between each ROI and its respective background. **(Shihan et al., 2021)**

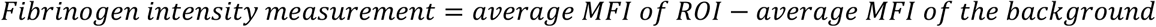

### Statistical analysis

All experiments were performed on at least three independent occasions (at least three biological replicates) and are indicated in the figure legends. Statistical analyses were performed via GraphPad Software, Inc., La Jolla, CA. Data was checked for normality using the Shapiro–Wilk or Kolmogorov–Smirnov Normality Test, and values are expressed as the means ± standard deviations (SDs) unless otherwise stated in the figure legends. To determine statistical significance, we selected appropriate tests for each examined parameter, including t tests for two-group comparisons and one- or two-way ANOVA for multigroup or multifactorial analyses. A suitable *post hoc* analysis was conducted to determine specific between-group differences for significant main effects or interactions identified through ANOVA. The results and corresponding figure legends provide comprehensive statistical details, including error representations, sample sizes, and experimental replication information. The results were considered statistically significant when the probability values (*p*) were less than or equal to the predetermined significance threshold of 0.05. A Pearson’s correlation was used to assess the strength and direction of the relationship between fibrinogen deposition and the time course of recruited macrophages in the noise-injured cochlea.

## RESULTS

### 1. Cre recombination efficiency in CX_3_CR1-lineage leukocytes in adult mouse cochlea and blood

We have previously employed *CX_3_CR1^GFP/wt^* reporter mice expressing enhanced green fluorescence protein (EGFP) in cochlear macrophages to study their distribution and function in the injured cochlea **(Kaur et al., 2019, KaurOhlemiller and Warchol, 2018, Kaur et al., 2015, Manickam et al., 2023)**. Although this mouse line aids in visualizing macrophages in the injured cochlea, it does not distinguish RM from recruited Mo/Mo-M. Therefore, it cannot be used to define the isolated roles of CX_3_CR1-RM vs. recruited Mo-M in SGN survival after injury. To address this, we have developed a fate mapping technique wherein CX_3_CR1-RM and recruited Mo/Mo-M are endogenously labeled with different fluorescent reporters. This was achieved by crossing mice bearing tamoxifen-inducible Cre (*CX_3_CR1^YFP−CreER/YFP−CreER^*) to a red fluorescence protein (RFP) reporter mouse line (*Rosa26^RFP^ or R26^RFP^*).(Yona et al., 2013, O’KorenMathew and Saban, 2016a, Plemel et al., 2020b) The progeny of this cross, *CX_3_CR1^YFP−CreER/wt^:R26^RFP^* mice were injected with either tamoxifen or corn oil (vehicle) at postnatal days 28 (P28) and 29 (P29) to label CX_3_CR1 expressing cells (i.e., YFP+ cell) with RFP and were euthanized at 2 (P32) and 60 (P90) days post injection (Fig. 1A). Our first aim was to determine the Cre recombination efficiency in CX_3_CR1 lineage in both blood and cochlea of an adult mouse by flow cytometry and immunohistochemistry, respectively. After 2 days of tamoxifen administration, we found 99% ± 0.39% of YFP+ RM in the cochlea and 40% ± 2.5% of YFP+ Mo in the blood expressed RFP. However, 60 days post-tamoxifen injection, only cochlear RM maintained RFP expression (Fig. 1B-J); whereas RFP expression in blood-circulating Mo was ‘washed out’ by 60 days, and only 2.5% ± 1.1% of these cells retained the expression (Fig. 1L, M). Thus, at the 60-day time-point, cochlear RM and blood-circulating Mo/Mo-M exhibited a YFP+ RFP+ and YFP+ RFP− phenotype, respectively. Tamoxifen injection did not affect the density of CD45-immunolabeled leukocytes in the cochlea, and the leukocyte density was comparable to that of mice injected with vehicle (Fig. 1K). We also found that in mice injected with vehicle (corn oil), 32% ± 3.3% of CX_3_CR1-YFP+ CD45+ RM in the cochlea and 1.9% ± 1.0% of CD45+ CD11b+ Ly6G-monocytes in the blood co-expressed RFP, which is indicative of a leaky Cre (Fig. 1B-E, M).

**Figure and figure legend 1.**
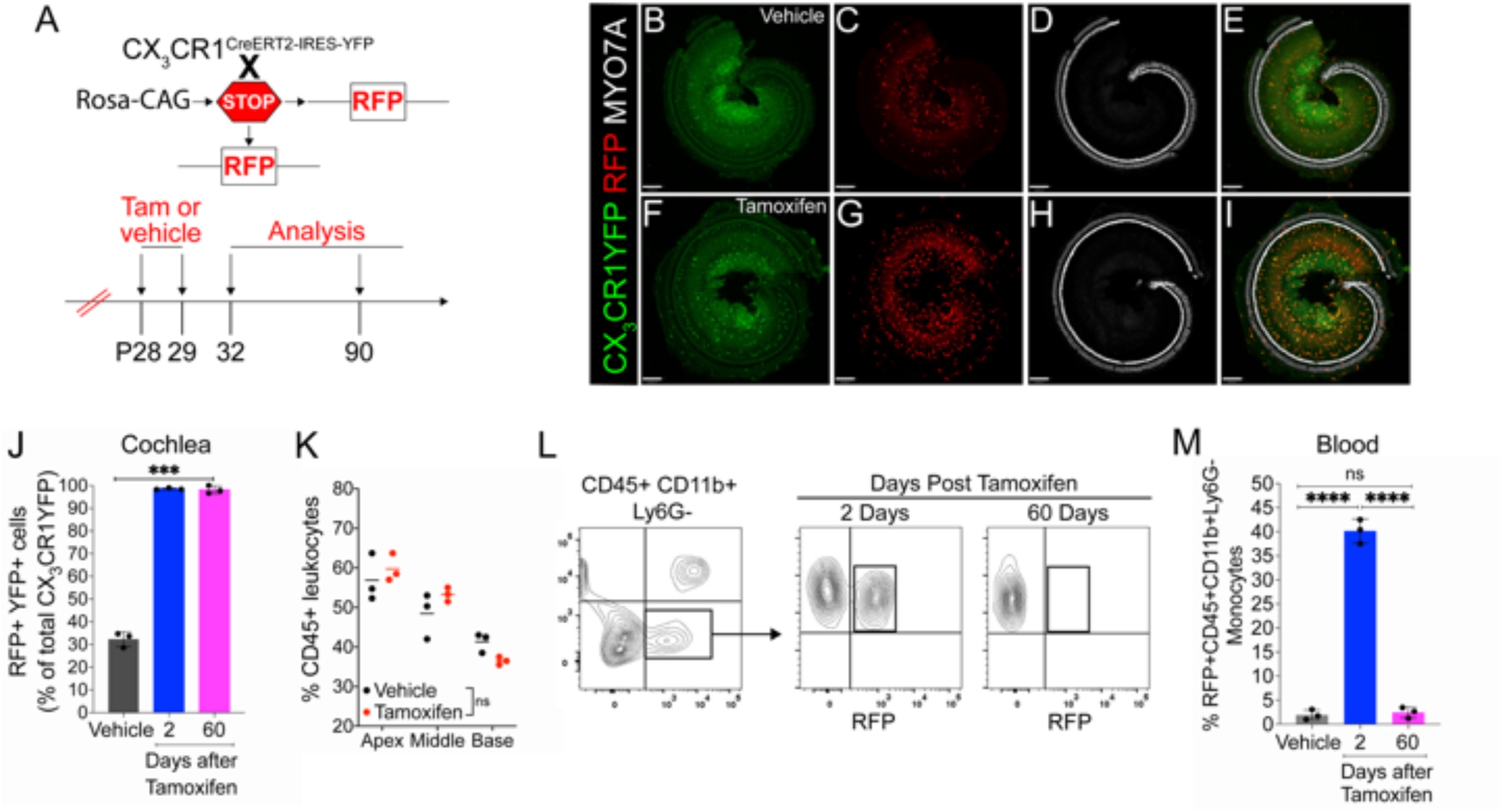
Cre recombination efficiency in CX_3_CR1-lineage leukocytes in adult mouse cochlea and blood in CX_3_CR1^YFP−CreER/wt^:R26^RFP^ mice. (A) Experimental regime. (B-I) Representative confocal images of cochlear whole mounts immunostained to label for B, F, CX_3_CR1-expressing macrophages (YFP/GFP, green), C, G Cre recombination (RFP/tdTomato, red), D, H, sensory hair cells (Myosin 7A, white), and E, I, merged (yellow) at 60 days post vehicle (corn oil) (B-E) or tamoxifen (F-I) injections. Scale bar = 130 µm. (J) Percentage of Cre recombination (RFP+) in YFP+ CX_3_CR1-expressing cochlear macrophages at 60 days after vehicle injections or at 2 and 60 days after tamoxifen injections. (K) Percentage of CD45+ leukocytes in the cochleae of vehicle- and tamoxifen-injected mice at 60 days post-injections. (L, M) Blood flow cytometry gating strategy (L) and quantification (M) of CX_3_CR1 lineage (CD45+, CD11b+, Ly6G-) (black box in L). N=3 mice per condition in J, K, and M. Data in J, K, and M are presented as Mean±SD. ***p<0.001, ****p<0.0001, ns, not significant; one-way ANOVA (J, M) or two-way ANOVA (K).

Our next aim was to determine the rate of turnover of cochlear RM in an adult mouse during steady state. To address this question, the progeny *CX_3_CR1^YFP−CreER/wt^:R26^RFP^* mice were injected with tamoxifen or corn oil and euthanized at various time points post injection (Fig. 2A). Examination of the recombined CX_3_CR1-expressing RM (i.e., YFP+ RFP+ cells) in different cochlear compartments such as the sensory epithelium, Rosenthal’s canal or spiral ganglion and lateral wall for more than one year indicate that their turnover rate is slower than the blood circulating Mo/Mo-M (typically ranging from 1 to 3 days) **(Gonzalez-Mejia and Doseff, 2009, WHITELAW and Bell, 1966)** but similar to the long-lived self-renewing microglia (brain resident macrophages) **(Ajami et al., 2007, Xu et al., 2007)** (Fig. 2B-F). Notably, a few recruited Mo/Mo-M (i.e., YFP+ RFP-cells) were observed under steady state, particularly in Rosenthal’s canal and spiral ligament as a function of aging, along with a trend of an increase in macrophage numbers in aged versus young cochlea (Fig. 2E, F).

**Figure and figure legend 2.**
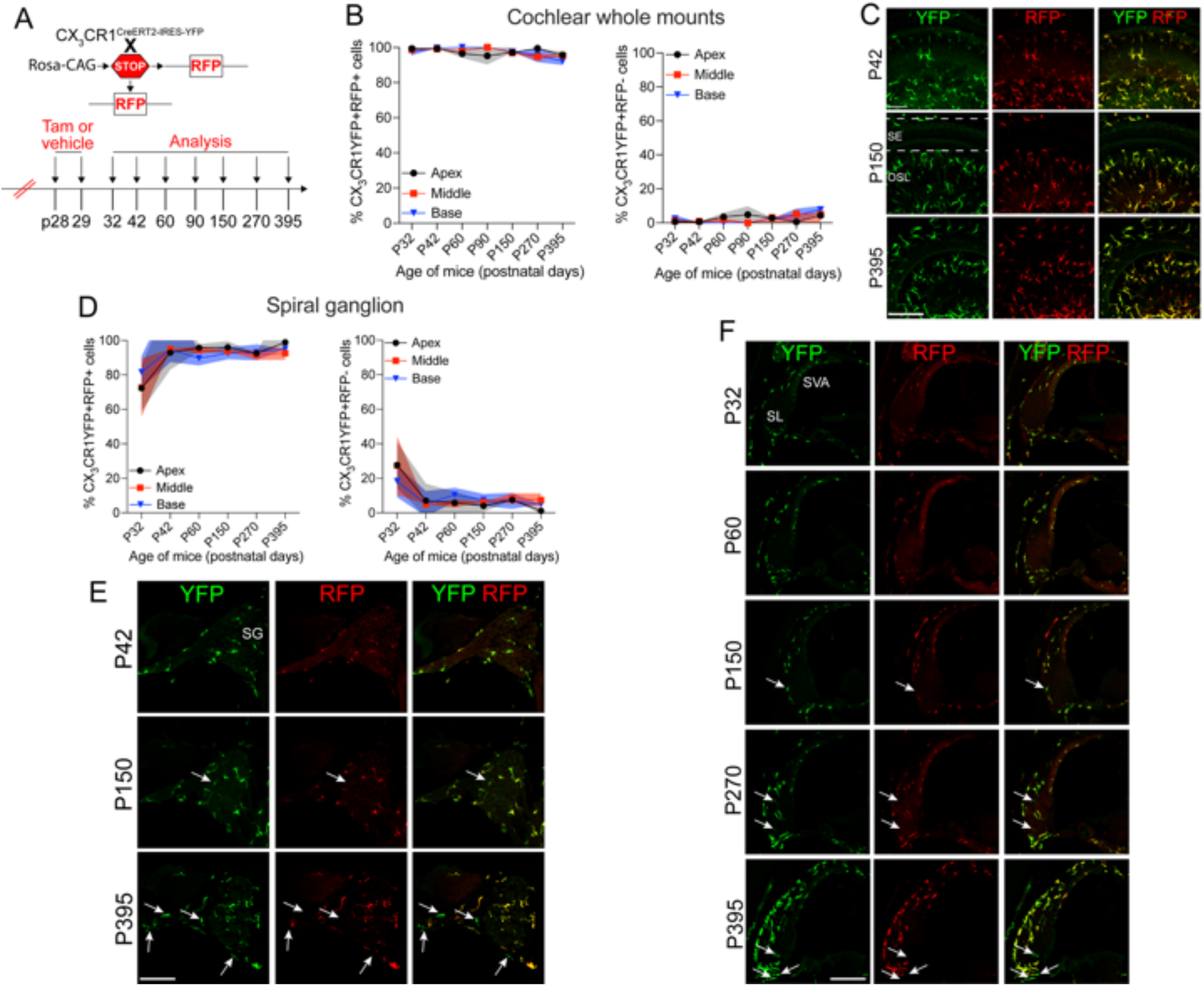
Slow turnover of cochlear resident macrophages in an adult mouse under steady state. (A) Experimental regime. (B) The percentage of CX_3_CR1YFP+ RFP+ RM (left) and CX_3_CR1YFP+ RFP-Mo/Mo-M (right) in mature cochlear whole mounts across a period of one year shows a slow turnover of resident macrophages in steady state. (C) Representative micrographs of the middle region of the cochlea show the presence of mostly the resident macrophages (YFP+ RFP+, yellow) at P42, P150, and P395. SE: sensory epithelium, OSL: Osseous spiral lamina. (D) The percentage of CX_3_CR1YFP+ RFP+ RM (left) and CX_3_CR1YFP+ RFP-Mo/Mo-M (right) in the spiral ganglion (SG) across a period of one year shows a slow turnover of resident macrophages in steady state. (E) Representative confocal micrographs of the spiral ganglion show the presence of mostly resident macrophages (YFP+ RFP+, yellow) and very few monocyte-derived macrophages (YFP+ RFP-, green, white arrows). (F) Representative confocal micrographs showing spiral ligament (SL) and stria vascularis (SVA) of the lateral wall of the basal cochlear turn show the presence of mostly resident macrophages (YFP+ RFP+, yellow) and very few monocyte-derived macrophages (YFP+ RFP-, green, white arrows). N=3 mice per time point in B and D. Data in B and D are presented as Mean±SD. Scale bar = 50 µm in all panels of C, E, and F. White arrows in E and F show the presence of YFP+ RFP-recruited macrophages (Mo/Mo-M) in the spiral ganglion and lower spiral ligament, respectively, as a function of biological aging.

### 2. Fate mapping with CX_3_CR1^YFP−CreER/wt^:R26^RFP^ mice distinguishes CX_3_CR1-expressing RM and recruited Mo/Mo-M in the cochlea following acoustic trauma

Previously, we have reported a unique spatiotemporal pattern of macrophage migration into the noise-injured cochlea, where macrophages migrate immediately and temporarily towards the damaged sensory epithelium, followed by a sustained increase in macrophage numbers in the spiral ganglion **(KaurOhlemiller and Warchol, 2018, Kaur et al., 2015)**. However, it is unclear whether this increase in macrophage numbers in the ganglion is due to local proliferation of RM or recruitment of Mo from blood circulation and their differentiation into Mo-M or both. To address this, we exposed the “washed out” mice to an acoustic trauma for 2 hours at a noise level of 112 dB SPL at 8-16 kHz that causes permanent hearing loss, and damage to hair cells and spiral ganglion neurons **(Hirose et al., 2005)**. Cochleae were analyzed immediately and at 1, 3, 5, 7, 10, 15, 30 days post acoustic trauma (Fig. 3A). We observed a YFP+ RFP+ population consistent with cochlear RM and a YFP+ RFP− population consistent with recruited Mo/Mo-M cells in the spiral ganglion by 7 days post exposure (Fig. 3B, bottom panel). In contrast, sham-exposed mice only contained YFP+ RFP+ RM (Fig. 3B, top panel). Next, we quantified the densities of YFP+ RFP+ RM and YFP+ RFP− recruited Mo/Mo-M in the neuronal region, i.e., spiral ganglion and spiral lamina, as a function of days post-exposure (DPNE) to determine their spatiotemporal distribution in the noise-injured cochlea. At 1 DPNE, the density of YFP+ RFP+ RM significantly decreased in the spiral ganglion and concurrently increased in the spiral lamina in the apex, middle, and basal cochlear turns when compared to densities in the sham-exposed and 0-day noise-exposed mice. Remarkably, by 3 DPNE, the density of YFP+ RFP+ RM in the spiral ganglion was restored to the levels as observed in the sham-exposed mice, and their numbers peaked by 5-7 DPNE and then returned to baseline by 30 DPNE in all cochlear turns. Similarly, the density of YFP+ RFP+ RM also peaked in the spiral lamina at around 3-7 DPNE in all three cochlear turns, after which they declined to the levels observed in the sham-exposed mice (Fig. 3C, D). YFP+ RFP− recruited Mo/Mo-M were rarely seen in and around the spiral ganglion and spiral lamina in the sham-exposed and 0-day noise-exposed cochlea. At 1 DPNE (i.e., same time when the numbers of YFP+ RFP+ RM are decreased in the spiral ganglion), YFP+ RFP− recruited Mo/Mo-M were found to be present in both spiral ganglion and spiral lamina. By 3-7 DPNE, the density of YFP+ RFP− recruited Mo/Mo-M increased in all three cochlear turns, with significantly higher densities in the apical ganglion and lamina, after which their numbers started to wane. By 30 DPNE, few (around 1-2) YFP+ RFP− recruited Mo/Mo-M were still observed in the spiral lamina but rarely in the spiral ganglion of all three cochlear turns (Fig. 3C, D). In addition to the neuronal region, Mo/Mo-M were also detected around the sensory epithelium and in the spiral ligament (primary site for macrophage accumulation), but not in the stria vascularis of the noise-injured cochlea (Fig. 4A-D).

**Figure and figure legend 3.**
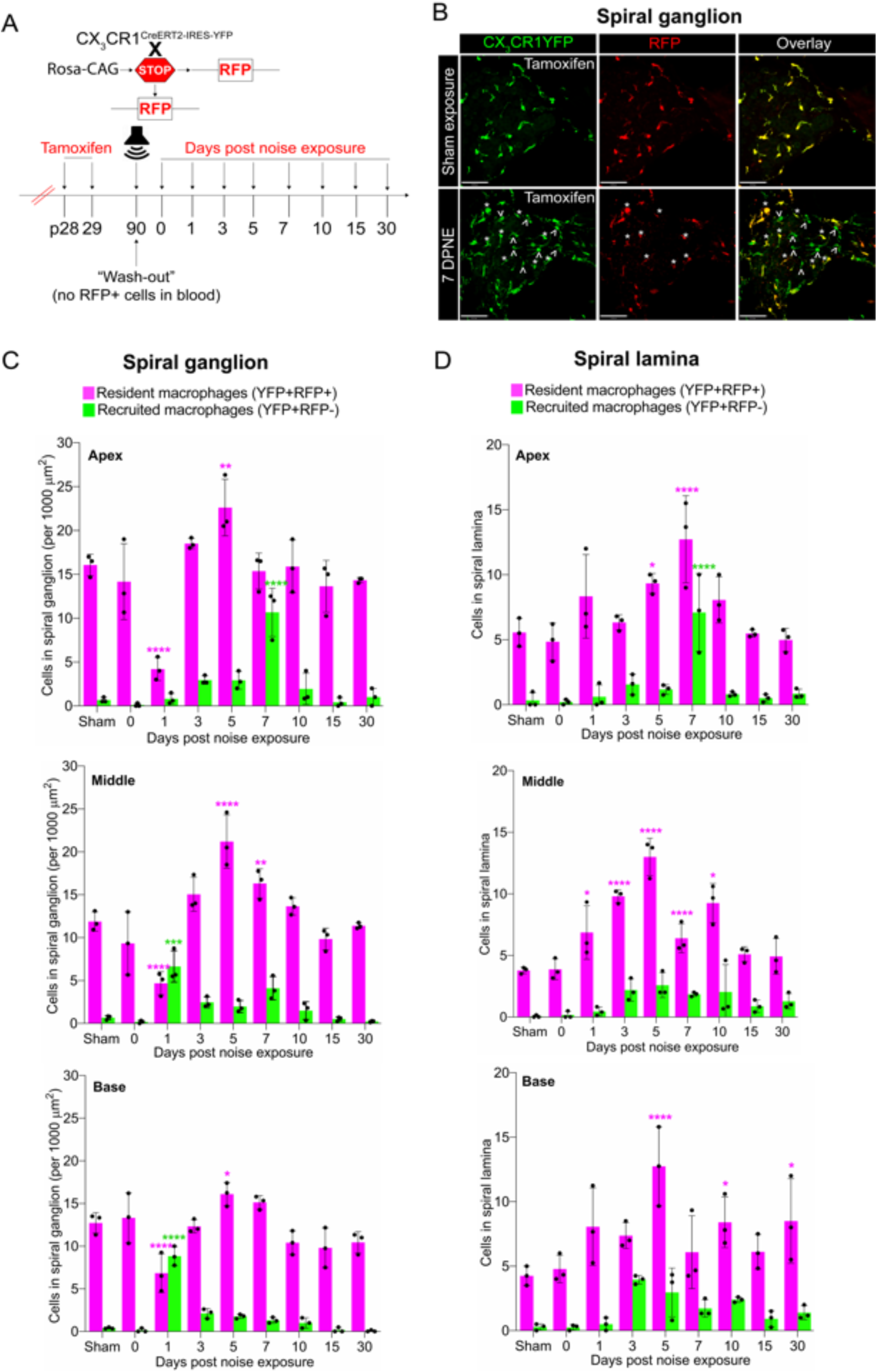
Fate mapping with CX_3_CR1^YFP−CreER/wt^:R26^RFP^ mice distinguishes CX_3_CR1-expressing RM and Mo/Mo-M in the neuronal region of the cochlea after acoustic trauma. (A) Experimental regime. (B) Representative confocal images of the spiral ganglion from tamoxifen-injected Cre mice after sham-exposure (top) and at 7 days after acoustic trauma (bottom) shows the presence of only resident macrophages (yellow overlay) after sham exposure and resident macrophages (YFP+ RFP+, yellow overlay, white asterisks) and recruited monocyte-derived macrophages (YFP+ RFP-, green overlay, white arrows) after acoustic trauma. Scale bar = 50 µm (C, D) Quantification of resident and recruited macrophages in the apical, middle, and basal spiral ganglion (C) and osseous spiral lamina (D) on different days after exposure. Data in C and D are presented as Mean±SD from N=3 mice per time point. *p<0.05, ** p< 0.01, ***p<0.001, ****p<0.0001; one-way ANOVA comparing macrophage numbers at different days post noise exposure with the sham group. Asterisks are color-coded based on resident (magenta) or recruited (green) macrophages, as shown in the graph legends.

**Figure and figure legend 4.**
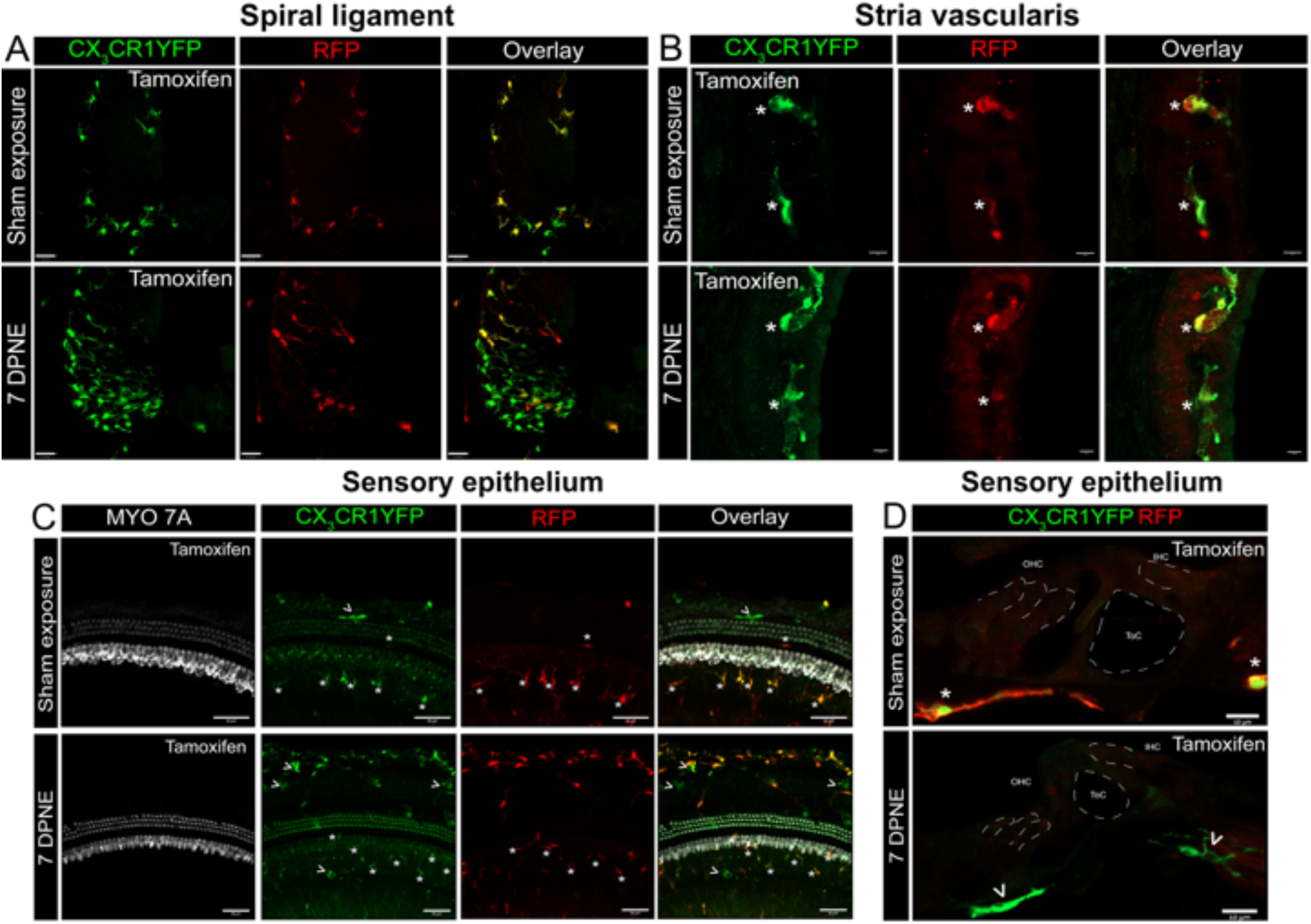
CX_3_CR1-expressing RM and Mo/Mo-M in the spiral ligament, stria vascularis, and sensory epithelium of the cochlea after acoustic trauma. Representative confocal images of the spiral ligament (A), stria vascularis (B), and sensory epithelium (C, D) from tamoxifen-injected Cre mice after sham-exposure (top) and at 7 days after acoustic trauma (bottom). White asterisks show YFP+ RFP+ resident macrophages (yellow overlay), and white arrows show YFP+ RFP-recruited monocyte/monocyte-derived macrophages (green overlay). Mo/Mo-M (green) are present in the lower spiral ligament as the primary site for macrophage accumulation (A), around the sensory epithelium (D), but not in the stria vascularis (B) of the noise-injured cochlea. Scale bars = 30 µm (A), 5 µm (B), 50 µm (C), and 10 µm (D).

### 3. Recruited Mo/Mo-M originate from bone marrow-derived circulating monocytes and are morphologically indistinguishable from cochlear RM undergoing *in situ* proliferation following acoustic trauma

Recruited macrophages into an injured tissue originate from bone marrow (BM) precursors, specifically, circulating monocytes that express high levels of Ly6C, CCR2, Ms4a3, and CD62L (L-selectin) and low levels of CX_3_CR1 **(CotechiniAtallah and Grossman, 2021, Gordon and Plüddemann, 2017, Liu et al., 2019, Yang et al., 2014)**. We immunolabeled the cochlear cryosections for the markers mentioned above to verify that the recruited Mo/Mo-M in the noise-injured cochlea originate from BM-derived circulating monocytes. Out of the four antibodies tested (Ly6C, CCR2, Ms4a3, and CD62L (L-selectin)), only two worked in our hands (Ly6C and CCR2). The data presented in Figure 5 show that the recruited Mo/Mo-M (YFP+ RFP−) expressed Ly6C and CCR2, confirming that these cells are derived from circulating monocytes (Fig. 5A, B). Unexpectedly, RM in the ganglion also displayed the expression of Ly6C (Fig. 5B) but were negative for CCR2 in the noise-injured cochlea.

**Figure and figure legend 5.**
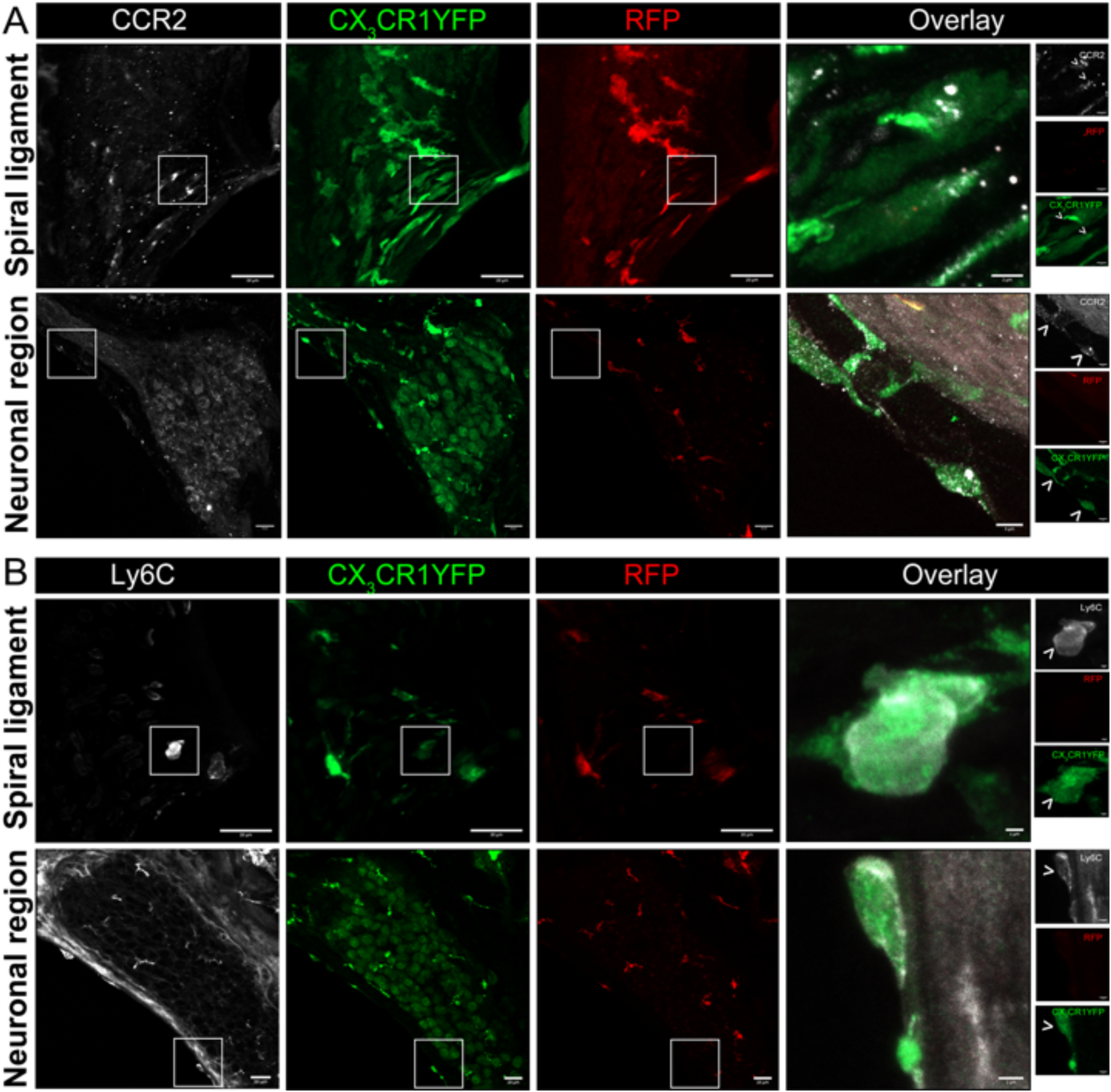
Blood-circulating recruited Mo/Mo-M express classical monocyte markers, CCR2 and LY6C, in the noise-injured cochlea. Representative confocal images showing expression of (A) CCR2 and (B) Ly6C in the blood-circulating recruited Mo/Mo-M (YFP+ RFP-) in the lower spiral ligament and neuronal region (Rosenthal’s canal and spiral lamina) after 3 days of exposure to 112 dB SPL noise level for 2 hours. Insets (white boxes) show the higher magnification of CCR2- and Ly6C-expressing recruited Mo/Mo-M in the overlay images. Note: Ly6C is also expressed in the RM (YFP+ RFP+) in the spiral ganglion at 3 DPNE (B), which could be associated with an inflammatory response/phenotype at the early stages of cochlear injury.

Morphometric analysis of YFP+ RFP+ RM and YFP+ RFP− Mo/Mo-M at 1- and 7-days post noise exposure displayed no significant morphological differences between the two cell types, suggesting that morphology is a poor indicator to distinguish CX_3_CR1-expressing resident and recruited macrophages in the noise-injured cochlea (Fig. 6A, B). Notably, recruited Mo transform from round or less ramified at 1 DPNE to being more ramified-like macrophages by 7 DPNE, suggesting that the recruited Mo may eventually differentiate into cochlear RM (Fig. 6A, B).

**Figure and figure legend 6.**
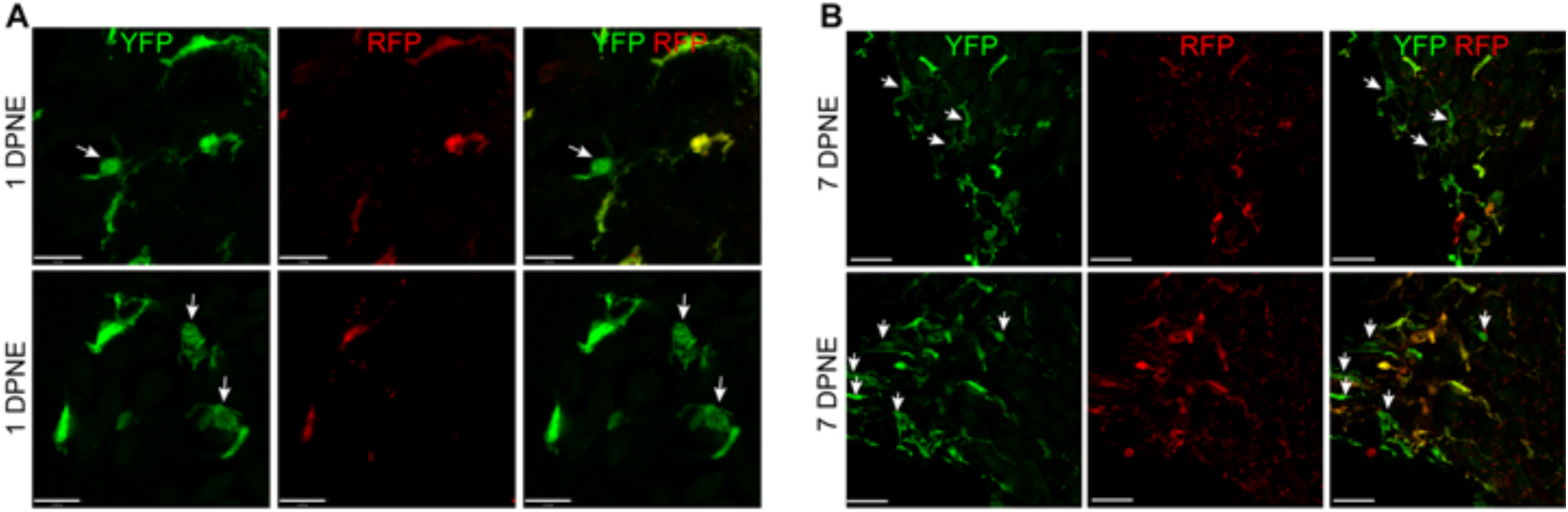
Morphology of CX_3_CR1-expressisng RM and Mo/Mo-M. Representative confocal micrographs showing indistinguishable morphology of CX_3_CR1-expressing resident (YFP+ RFP+) and recruited (YFP+ RFP-, white arrows) macrophages in the spiral ganglion at 1 (A) and 7 (B) days post-noise exposure. Note: The recruited circulating Mo was transformed from being round or less ramified at 1 DPNE to more ramified by 7 DPNE (white arrows). This suggests that the infiltrated circulating Mo may differentiate into cochlear RM. Scale bars = 17 µm (top panel), 13 µm (bottom panel) in A and 20 µm in B, respectively.

Microglia (brain RM) undergo local proliferation after injury **(Plemel et al., 2020a)**. Thus, to determine if acoustic trauma induces RM expansion in the spiral ganglion (as seen in Fig. 3) due to local proliferation and limited death, cochleae were immunolabeled for Ki-67 (cell proliferation marker) and cleaved caspase-3 (CC3, key effector of apoptosis), and the numbers of Ki67+ and CC3+ RM and Mo/Mo-M were quantified. Probing for cleaved caspase-3 showed no evidence for RM and Mo/Mo-M undergoing apoptosis (Fig. 7A). As expected, immunolabeling for Ki67 showed proliferating YFP+ RFP− Mo/Mo-M only at 1 DPNE (i.e., time when these cells infiltrate the spiral ganglion) in the middle and basal spiral ganglion as they typically proliferate during the process of extravasation from the blood into the noise-injured cochlea (Fig. 7B, C). Unexpectedly, Ki67 immunolabeled YFP+ RFP+ RM were observed at 3 DPNE in the spiral ganglion of all three turns, indicating that RM undergo *in-situ* proliferation after acoustic trauma (Fig. 7B, C). These and the above data imply that the increase in overall macrophage numbers in the spiral ganglion and lamina after acoustic trauma results from both *in situ* proliferation of RM and recruitment of blood-circulating Mo/Mo-M. Although blood-derived macrophages acutely infiltrated the spiral ganglion, resident macrophages progressively monopolized the ganglion after noise trauma, likely due to proliferation.

**Figure and figure legend 7.**
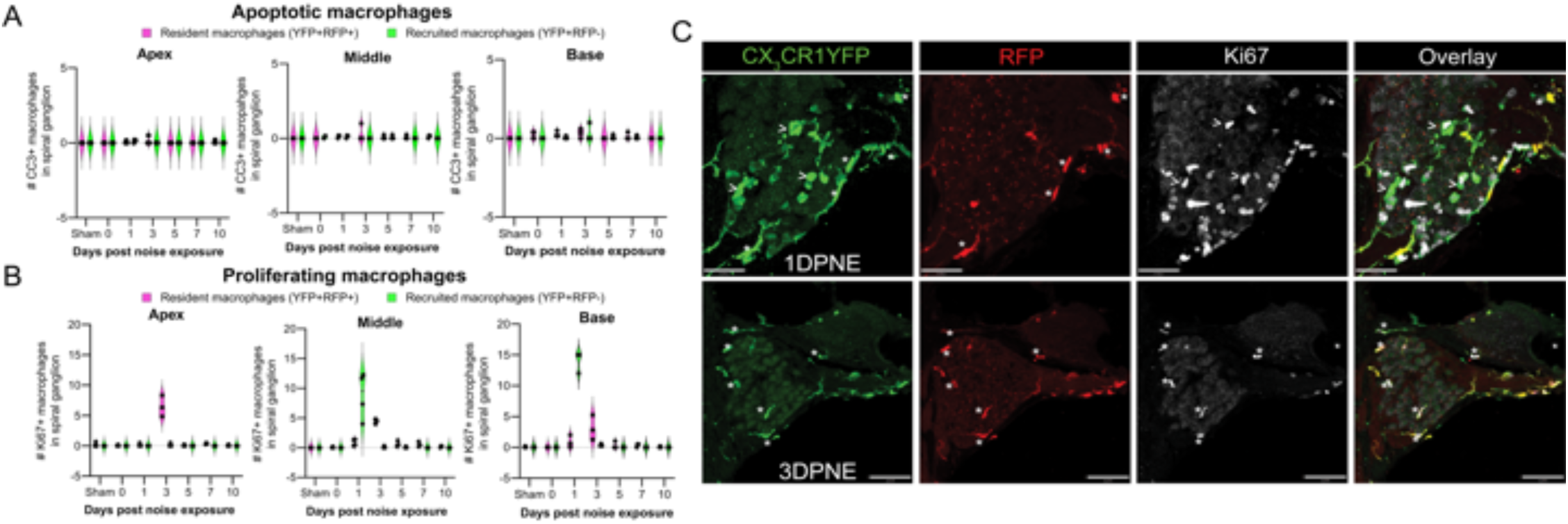
Fate of CX_3_CR1-expressisng RM and Mo/Mo-M. Number of (A) Cleaved caspase 3 (CC3) and (B) Ki67 positive macrophages in the apical, middle, and basal spiral ganglion on different days after noise exposure. Data is plotted as Mean±SD. N=3-4 mice per recovery time. (C) Representative confocal micrographs showing Ki67 positive resident (YFP+ RFP+, white asterisks) and recruited (YFP+ RFP-, white arrows) macrophages in the spiral ganglion of the middle turn at 1- and 3-days post noise exposure. Scale bar = 31 µm (top panel) and 70 µm (bottom panel) in C.

### 4. Recruitment of CX_3_CR1-expressing Mo/Mo-M into the spiral ganglion is associated with vascular damage and fibrinogen extravasation from blood into the cochlea after acoustic trauma

The blood labyrinth barrier (BLB) is a network of specialized capillaries that normally limits the passage of substances, including toxins, proteins, pathogens, and immune cells, between the bloodstream and inner ear fluids, which is crucial for the maintenance of cochlear homeostasis. Inflammation, noise trauma, ototoxicity, and aging produce changes in the BLB, resulting in increased vascular permeability, also known as “leaky vasculature”**(Hirose et al., 2014, Shi, 2016)**. Such leaky vasculature may allow the blood-circulating immune cells to enter the compromised cochlea. Therefore, we wanted to determine if recruitment of CX_3_CR1-expressing Mo/Mo-M in the neuronal region occurs either before, at the same time, or after the breakdown of the BLB in the noise-injured cochlea. To address this, we immunolabeled the cochlear sections from the sham-exposed and noise-exposed “washed out” mice with fibrinogen antibody. Fibrinogen is a pleiotropic blood-clotting protein that extravasates into the nervous system after injury or disease associated with vascular damage or blood-brain barrier (BBB) breakdown, thus serving as a marker of BBB disruption **(RyuDavalos and Akassoglou, 2009)**. The data shows fibrinogen immunolabeling inside the cochlear labyrinth after an acoustic trauma of 112 dB SPL for 2 hours at 8-16 kHz octave band, suggesting vascular damage or compromised BLB following such injury (Fig. 8A). Fibrinogen extravasation and deposition was observed inside and around the spiral ganglion, in the spiral limbus, as well as in the lower spiral ligament and stria vascularis, which are predominantly vascularized cochlear regions, suggesting vascular damage after exposure (Fig. 8A). Compared to the uninjured cochlea, the fluorescence intensity of fibrinogen increased by 1-3 DPNE primarily in the middle and basal spiral ganglion and spiral limbus, and in the lower spiral ligament and was decreased by 10 DPNE (Fig. 8A, C). To determine if the recruitment of blood-circulating Mo/Mo-M in the SGN is associated with a leaky vasculature after acoustic trauma, a correlation analysis was performed between fibrinogen intensity and recruited Mo/Mo-M density in the spiral ganglion at 0, 1, 3, 5, 7, and 10 days after noise exposure. The analysis revealed a strong and positive correlation between fibrinogen intensity and macrophage recruitment in the middle ganglion (Pearson correlation coefficient, r = 0.8), whereas no correlation was found in the base and apex ganglia of the noise-injured cochlea (r = 0.16 and -0.09, base and apex, respectively) (Fig. 8D). These data offer a novel probe, fibrinogen, as a blood marker for cochlear vascular damage after acoustic trauma and indicate that recruitment of circulating Mo/Mo-M in the cochlea follows vascular damage.

**Figure and figure legend 8.**
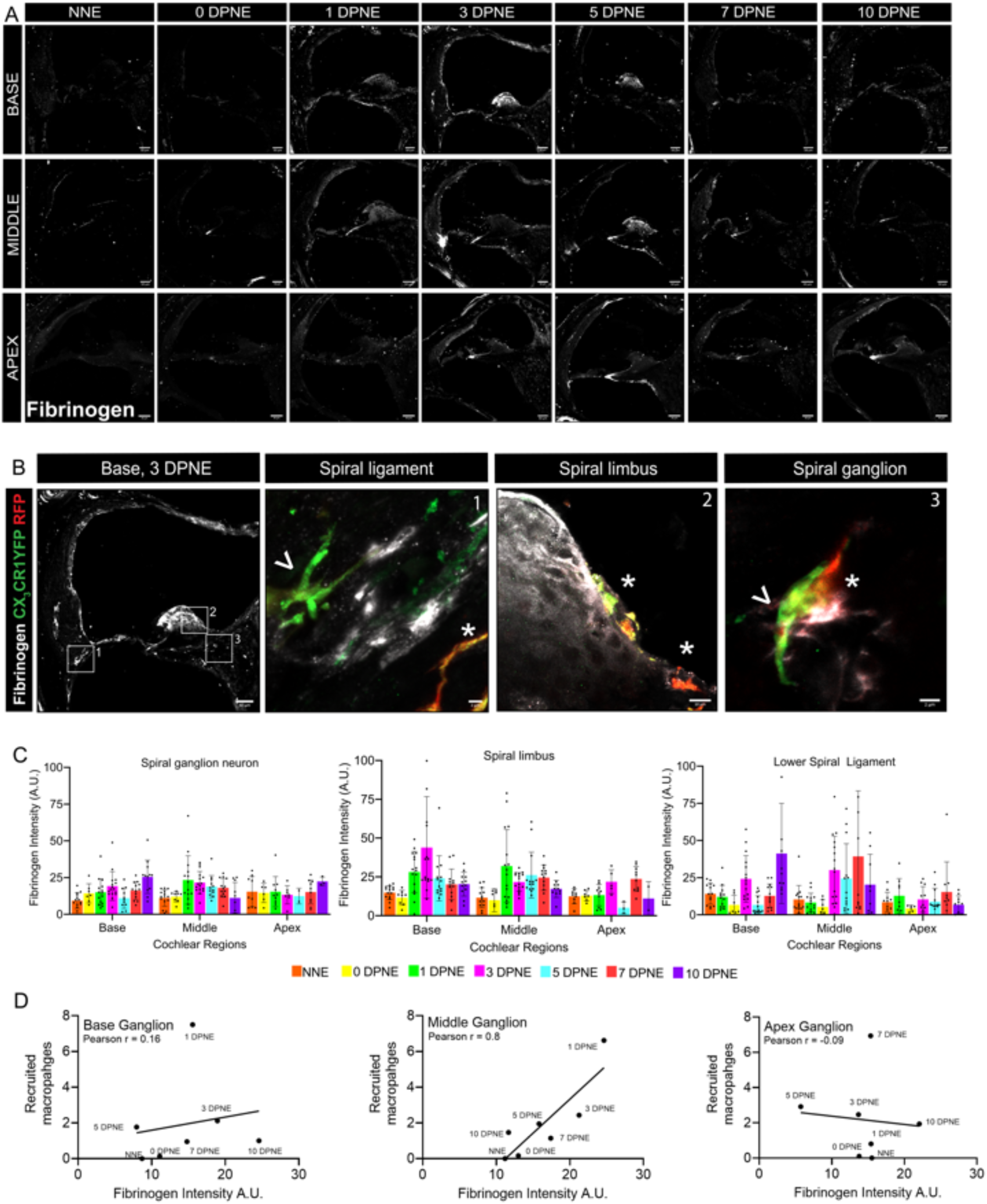
Fibrinogen extravasation from blood and deposition in the cochlear labyrinth indicates vascular aberration and positively correlates with the recruitment of Mo/Mo-M after acoustic trauma. (A) Representative confocal images of fibrinogen extravasation in the base, middle, and apex cochlear turns at different time points after an acoustic trauma of 112 dB SPL for 2 hours at 8-16 kHz octave band. Scale bar = 50 µm. (B) Representative high magnification confocal images showing fibrinogen fluorescence and juxtaposed RM (white asterisk) and Mo/Mo-M (white arrow) in the lower spiral ligament (1), spiral limbus (2), and spiral ganglion (3) at 3 DPNE. (C) Fibrinogen fluorescent intensity in arbitrary units (A.U.) in apex, middle, and base spiral ganglion, lower spiral ligament, and spiral lamina as a function of days post noise exposure (DPNE). Data is plotted as Mean±SD. Each dot in the graphs in C represents cochlear cross sections from 2 to 4 animals per time point. (D) Pearson’s correlation analysis between fibrinogen intensity and recruited Mo/Mo-M density in the base, middle, and apex spiral ganglia at different time points after an acoustic trauma. r, Pearson correlation coefficient.

## DISCUSSION

Our novel work involves the use of fate mapping with *CX_3_CR1^YFP−CreER/wt^:R26^RFP^* mice to address unresolved questions concerning the presence of bona fide resident versus recruited macrophages and their phenotypic differences and similarities in the injured cochlea. Distinguishing resident from recruited macrophages in the injured cochlea has proven to be challenging. Indeed, recruited classical Mo can be resolved phenotypically acutely after injury as they express Ly6C and CCR2, but not F4/80 or Iba-1 **(Pan et al., 2024, Shin et al., 2022)** . However, differentiated Mo-derived macrophages (Mo-M) may overlap phenotypically with RM because both express CX_3_CR1, F4/80, or Iba-1, but non- or reduced expression of Ly6C **(Shin et al., 2022)**. Using genetic reporter mice such as *CX_3_CR1^GFP/wt^* mice to identify CX_3_CR1-expressing RM in the injured cochlea can be misleading, because recruited Mo/Mo-M also express CX_3_CR1 **(Jung et al., 2000, Molawi et al., 2014, Yona et al., 2013, Zigmond et al., 2012)**, as shown in the central nervous system (CNS) **(Kierdorf et al., 2013, Shechter et al., 2013)**. A similar concern is also with using *CCR2^RFP/wt^* mice to identify Mo and Mo-derived cells, as fully differentiated Mo-M downregulate their expression of CCR2 to negligible levels that are similar to RM, such as microglia in the brain **(GreterLelios and Croxford, 2015, Saederup et al., 2010)**. Generating GFP bone marrow chimeras is a commonly used non-genetic approach to discern cochlear RM from recruited Mo and Mo-derived cells under steady state, or during injury **(Lang et al., 2016, Okano et al., 2008, Sato et al., 2008, Shi, 2010)**. However, this method is also confounded since whole-body irradiation/bone marrow transplantation itself leads to recruitment of bone marrow-derived cells into the tissue **(Ajami et al., 2007, Chen et al., 2012, Okano et al., 2008, Xu et al., 2007)**. Parabiosis bypasses this problem **(Ajami et al., 2007)** but does not reach full chimerism in the periphery and is not widely used for multiple reasons **(Kierdorf et al., 2013)**, including significant technical challenges.

In the current study, we found that the *CX_3_CR1^YFP−CreER/wt^:R26^RFP^* mouse induces *loxP* recombination in cochlear RM (∼ 30%) and blood myeloid cells (∼ 2%) in the absence of tamoxifen. Such *CreER* leakiness has also been reported in microglia of *CX_3_CR1^CreER^* mouse lines **(Fonseca et al., 2017, Stowell et al., 2019, Van Hove et al., 2020)**. Nevertheless, this *CreER* leakiness did not significantly hinder the interpretation of the data regarding cochlear RM turnover under steady state or recruitment, distribution, and fate of RM, and Mo/Mo-M in the aging or noise-injured cochlea. This is because of the use of the ‘washed out’ *CX_3_CR1^YFP−CreER/wt^:R26^RFP^*mice in which, upon tamoxifen injection, nearly 100% of cochlear RM recombined, whereas blood myeloid cells lose the recombination by 60 days, as they are short-lived and turn over quickly due to ongoing, lifelong hematopoiesis. Besides *CX_3_CR1*, cochlear RM also expresses *Tmem119* and *P2ry12* **(Bassiouni et al., 2023, Chiot et al., 2024)**, and inducible *Cre* lines, including *Tmem119^CreER^* and *P2ry12^CreER^* are available commercially. A comparative analysis of these inducible lines for *Cre* recombination with and without tamoxifen, as well as specificity (resident vs. recruited), is needed in the future to inform on the caveats and benefits of all tools for reliable manipulation of cochlear macrophage function *in vivo* and development of macrophage-based therapies for acquired sensorineural hearing loss. Studies using bone marrow chimeras have reported that the turnover of cochlear RM is slow and progressive under steady state **(Okano et al., 2008, Shi, 2010)**. Our findings with the *CX_3_CR1^YFP−CreER/wt^:R26^RFP^*fate mapping tool corroborate with these studies. They reveal that while RM represented a stable population in the cochlea, a slow and progressive contribution of blood-circulating Mo/Mo-M to the macrophage pool occurs in the spiral ganglion and lateral wall, starting around 5 months of age. With respect to turnover rates, our findings imply that cochlear RM have characteristics as of resident macrophages/microglia in the brain and retina that are long-lived **(Ajami et al., 2007, Hess et al., 2004, Simard et al., 2006, Xu et al., 2007)** and not as peripheral tissue-resident macrophages that turnover quickly **(Gonzalez-Mejia and Doseff, 2009, Naito et al., 1997)**. The mechanisms that regulate the maintenance of cochlear RM remain to be elucidated and are subject to future investigation.

Our data with acoustic trauma in the ‘washed out’ *CX_3_CR1^YFP−CreER/wt^:R26^RFP^* mice revealed interesting spatial and temporal distribution patterns for cochlear RM and Mo/Mo-M, particularly in the neuronal region. A day after acoustic trauma, the decreased density of RM in the spiral ganglion and their simultaneous increase in the spiral lamina across all cochlear turns validate our previous work with a 2-hour exposure at 120 dB SPL reported in Kaur et al., 2018. Reduced number of RM in the spiral ganglion is not due to apoptosis and may suggest that RM in the spiral ganglion migrate towards the spiral lamina and sensory epithelium immediately following acoustic trauma. During the same time after exposure, Mo/Mo-M infiltrate the empty niche in the middle and base spiral ganglion. The molecular signals that allow local migration of RM from the ganglion towards the lamina and that cause Mo/Mo-M extravasation from the blood into the ganglion remain unknown. We have previously reported the fractalkine (CX_3_CL1) molecule as a putative chemoattractant for macrophages that also regulate their numbers in the injured cochlea **(KaurOhlemiller and Warchol, 2018, Kaur et al., 2015, Manickam et al., 2024)**. Additional studies are needed to determine if fractalkine is a chemoattractant for RM or Mo/Mo-M or both in the injured cochlea. Unexpectedly, the reduced numbers of RM (YFP+RFP+) in the spiral ganglion started to recover by 3 days and were restored to baseline by 1 week following acoustic trauma. One reason for this effect would be the *in situ* proliferation of the remaining cochlear RM. To this end, our data establishes for the first time the presence of Ki67-positive cochlear RM (YFP+RFP+) at 3 days post-noise exposure, which may contribute to the pool of increased numbers of RM. Another explanation could be that recruited Mo (YFP+ RFP-) in the ganglion immediately and 1 day after exposure quickly differentiate into RM, and thus increase the density of RM. Indeed, **(Shin et al., 2022)** have reported that after acoustic overstimulation, monocytes infiltrated the lower spiral ligament within 2 days, followed by transformation into macrophages at 3-5 days based on CX_3_CR1 upregulation and Ly6C downregulation immunolabeling. Nonetheless, in the fate mapping ‘washed out’ *CX_3_CR1^YFP−CreER/wt^:R26^RFP^* mice, recruited Mo/Mo-M (YFP+ RFP-) in the spiral ganglion or lower spiral ligament did not show the expression for RFP, in case they differentiated into RM. While our data show that the recruited Mo/Mo-M expressed classical monocyte markers, Ly6C and CCR2, their further characterization for expression of unique markers for RM, including Iba-1, F4/80, CD64, and MHC-II, is vital to resolve this issue. Besides recruited Mo/Mo-M (YFP+ RFP-), Ly6C expression was also observed in RM (YFP+RFP+) in the spiral ganglion at 3 days post-noise exposure. Such Ly6C expression on both resident and recruited macrophages could be associated with an inflammatory response or phenotype during the early stages of cochlear injury. Whether these Ly6C-expressing RM switch to non- or low-Ly6C expression at later stages of injury remains to be determined. **(Hirose et al., 2005)** reported that an increase in macrophages in the noise-injured cochlea arises from infiltration of inflammatory cells from the vasculature as opposed to mitotic division of resident phagocytes. Yet, our findings using the fate mapping across different days after acoustic trauma have shed new light on the basic understanding of the mechanisms of the increase in macrophage numbers in the spiral ganglion in the noise-injured cochlea, which involves both recruitment of blood-circulating Mo and *in situ* proliferation of cochlear RM. Fibrinogen, also known as coagulation factor I, is abundant in the blood and functions in the coagulation cascade, creating polymers of fibrin. Fibrinogen leakage into the brain or retina is a hallmark of a compromised blood-brain or blood-retina barrier in multiple sclerosis **(DavalosMahajan and Trapp, 2019)**, Alzheimer’s **(Ryu and McLarnon, 2008),** and diabetic retinopathy patients **(MurataIshibashi and Inomata, 1992)**. Similarly, it has been reported that there is a strong relationship between plasma fibrinogen levels and the pathogenesis of sudden sensorineural hearing loss **(Lu et al., 2008)**. In this study, we show that fibrinogen extravasation and deposition occur in the cochlea as early as one day after acoustic trauma in areas closely associated with blood vessels, including the spiral ligament, the spiral limbus, and the spiral ganglion. Other investigators have also shown leaky cochlear vessels with fluorescent tracers such as serum albumin-FITC or serum protein IgG in noise-exposed animals **(Yang et al., 2011, Zhang et al., 2013)**. However, fibrinogen is a large glycoprotein with a molecular weight of 340 kDa relative to 65 kDa of albumin or 150 kDa of IgG, highlighting the magnitude of BLB damage after a 2-hour exposure to a noise level of 112 dB SPL. Remarkably, such fibrinogen deposition strongly correlated with the time course of recruitment of blood-circulating Mo/Mo-M primarily in the spiral ganglion and with perivascular clustering of activated macrophages in the ligament, limbus, and spiral ganglion of the noise-injured cochlea. This suggests that the dynamics of recruitment of circulating Mo/Mo-M is determined by leaky vasculature. Studies have revealed structural and molecular changes in the BLB after acoustic trauma that result in destabilization of the barrier and thus cause leaky vasculature. These changes include alterations in the location of pericytes, activation of perivascular macrophages, production of PEDF by activated macrophages that results in downregulation of barrier tight-junction and adherent-junction associated proteins, and vascular leakage **(Shi, 2009, Yang et al., 2011, Zhang et al., 2013, Zhang et al., 2012)**. The process by which blood-circulating monocytes are recruited into an injured tissue is called chemotaxis, which is the directed movement of cells in response to chemokines. Several adhesion molecules and chemoattractants, including ICAM-1, MCP-5 (CCL12), MCP-1 (CCL2), and MIP-1beta (CCL4), have been found to be upregulated after acoustic trauma **(Frye et al., 2017, Sautter et al., 2006, TanThorne and Vlajkovic, 2016, Tornabene et al., 2006)**. Chemokine CCL2 and its primary receptor, CCR2, are the most widely validated effector of monocyte chemotaxis *in vivo* **(Ransohoff et al., 2002, Rollins, 1996)**. However, it has been shown that the number of cochlear macrophages remains unchanged after acoustic trauma in the absence of CCL2 or CCR2 **(Sautter et al., 2006).** Investigations on the specific chemoattractants that recruit Mo/Mo-M into the noise-injured cochlea and determining their molecular phenotypes and precise functions after acoustic trauma are underway.

In conclusion, this study demonstrates for the first time the use of fate-mapping as a tool to clearly distinguish cochlear resident macrophages from blood-circulating recruited monocytes and monocyte-derived macrophages in normal, aging, and noise-injured cochleae. Our data confirms that cochlear RM turnover occurs at a slower rate than the peripheral leukocytes. In addition, biological aging or acoustic trauma recruit blood-circulating Mo into the cochlea through a leaky/permeable BLB, which are morphologically similar to RM. Importantly, besides the recruitment of Mo and their differentiation into tissue macrophages (Mo-M), self-renewal of RM also contributes to the overall increase in the numbers of macrophages in the cochlea after acoustic trauma (Fig. 9).

**Figure and figure legend 9.**
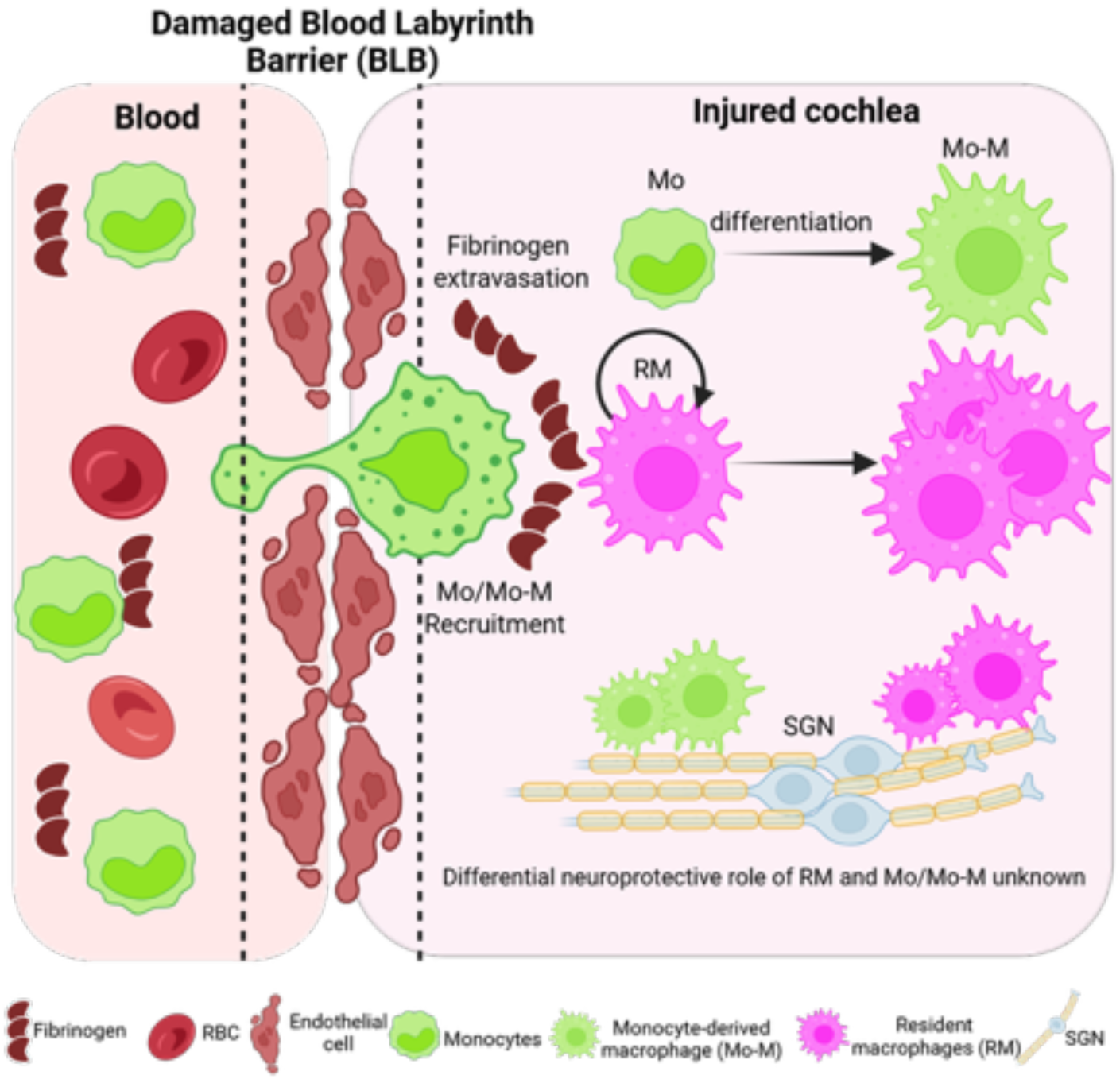
Working model of the ontogeny and dynamics of resident and recruited macrophages in the injured cochlea. Schematic illustrating disruption of the blood-labyrinth barrier (BLB), fibrinogen glycoprotein extravasation from blood, and subsequent recruitment of circulating monocytes (green) into the injured cochlea following an acute (2 hours) acoustic trauma imparting permanent hearing loss. The short-lived recruited monocytes may ultimately differentiate into macrophages (monocyte-derived macrophages, Mo-M). The long-lived cochlear resident macrophages (RM, magenta) undergo self-renewal and expansion in the injured cochlea. The overall increase in the number of macrophages in the acute noise-injured cochlea is attributed to the recruitment of blood-circulating monocytes, their prospective differentiation into macrophages, and the *in-situ* proliferation of resident (local) macrophages. Defining the dynamics and the differential neuroprotective functions of CX_3_CR1-expressing cochlear resident and recruited macrophages in acute and chronic pathological cochleae is under investigation. The illustration is created in BioRender.

## abbreviations

RM: Resident macrophages
Mo: Monocytes
Mo-M: Monocyte-derived macrophages
YFP: Yellow fluorescent protein
GFP: Green fluorescent protein
RFP: Red fluorescent protein
IHC: Inner hair cell
SGN: Spiral ganglion neuron
OSL: Osseous spiral lamina
SE: Sensory epithelium
dB: Decibel
SPL: Sound pressure level
kHz: Kilohertz
kDa: Kilodalton
DPNE: Days post noise exposure
BLB: Blood labyrinth barrier

## DECLARATIONS

### Availability of data and materials

All data are available in the main text or the supplementary materials.

## Acknowledgements

We thank Dr. Astrid Cardona and her lab members at the University of Texas, San Antonio (UTSA) for sharing the flow cytometry protocol for immunophenotyping of CX_3_CR1 lineage in the blood of adult mice.

## Author’s contributions

TK conceptualized and designed the study. SVM, ARS, LB, and TK ran experiments. SVM, ARS, EP, and TK analyzed and interpreted data. TK and SVM co-wrote the manuscript.

## Ethics declarations

### Ethics approval and consent to participate

All aspects of animal care, procedure, and treatment were carried out according to the guidelines of the Animal Studies Committee of Creighton University, Omaha, NE, and Rutgers University, Piscataway, NJ.

